# Targetable genetic alterations of *TCF4* (E2-2) drive immunoglobulin expression in the activated B-cell subtype of diffuse large B-cell lymphoma

**DOI:** 10.1101/170605

**Authors:** Neeraj Jain, Keenan Hartert, Saber Tadros, Warren Fiskus, Ondrej Havranek, Man-Chun John Ma, Alyssa Bouska, Tayla Heavican, Dhiraj Kumar, Qing Deng, Dalia Moore, Christine Pak, Chih Long Liu, Andrew Gentles, Elena Hartmann, Robert Kridel, Karin Ekstrom Smedby, Gunnar Juliusson, Richard Rosenquist, Randy D. Gascoyne, Andreas Rosenwald, Filippo Giancotti, Sattva S. Neelapu, Jason Westin, Julie Vose, Matthew Lunning, Timothy Greiner, Scott Rodig, Javeed Iqbal, Ash Alizadeh, R. Eric Davis, Kapil Bhalla, Michael R. Green

## Abstract

The activated B-cell (ABC) subtype of diffuse large B-cell lymphoma (DLBCL) is characterized by the chronic activation of signaling initiated by immunoglobulin-μ (IgM). By analyzing DNA copy profiles of 1,000 DLBCLs, we identified gains of 18q21.2 as the most frequent genetic alteration in ABC-like DLBCL. We show that these alterations target the *TCF4* (E2-2) transcription factor, and that over-expression of *TCF4* leads to its occupancy on immunoglobulin gene enhancers and increased expression of IgM at the transcript and protein level. The *TCF4* gene is one of the top BRD4-regulated genes in DLBCL. Using a BET proteolysis-targeting chimera (PROTAC) we show that TCF4 and IgM expression can be extinguished, and ABC-like DLBCL cells can be killed *in vitro* and *in vivo*. This highlights a novel genetic mechanism for promoting immunoglobulin signaling in ABC-like DLBCL and provides a functional rationale for the use of BET inhibitors in this disease.

## INTRODUCTION

Diffuse large B-cell lymphoma (DLBCL) is the most common form of lymphoma and is curable in ∼60% of patients using a combination chemo-immunotherapy regimen, R-CHOP^1,2^. However, those that are refractory to, or relapse following, first-line therapy have a dismal outcome^3^. Chimeric antigen receptor (CAR)-T cells are likely to change the landscape of outcomes in relapsed/refractory patients, but a large number of patients are not eligible for CAR-T therapy and ∼50% of those that received CAR-T progress within 12 months^4^. Novel rationally-targeted therapeutic strategies are therefore needed for DLBCL.

The clinical heterogeneity of DLBCL is underpinned by molecular heterogeneity, with the major distinction being between the germinal center B-cell (GCB)-like and activated B-cell (ABC)-like ‘cell of origin’ (COO) subtypes that were identified by gene expression profiling^5^. The GCB-like subtype shows transcriptional similarities to normal germinal center B-cells, whereas the ABC-like subtype shows transcriptional similarities to CD40-activated B-cells or plasmablasts. Patients with ABC-like DLBCL have significantly worse overall survival compared to patients with GCB-like DLBCL, when treated with the standard-of-care combination chemotherapy (CHOP) plus rituximab (R-CHOP) regimen^6^. The ABC-like DLBCL subtype expresses immunoglobulin μ (IgM)^7^ in >90% of cases, which forms the B-cell receptor (BCR) signaling complex in association with CD79A and CD79B and drives chronically active BCR signaling. Several genetic alterations have been shown to promote this signaling, including mutations of the *CD79A, CD79B, CARD11*, and *MYD88* genes^8-11^. However, these mutations only account for approximately two thirds of ABC-like DLBCL cases^12^, suggesting that other significant genetic drivers remain to be defined.

A common mechanism for tumorigenesis is the gain or loss of DNA encoding oncogenes or tumor suppressor genes, respectively. These copy number alterations (CNAs) perturb a higher fraction of the cancer genome than somatic nucleotide variants (SNVs) and small insertion/deletions (InDels) and are critically important to cancer etiology^13^. Here, we have integrated multiple datasets, including DNA copy number profiles of 1,000 DLBCLs, and identified DNA copy number gain of the E2 transcription factor *TCF4* as the most frequent genetic alteration in ABC-like DLBCL. We show that TCF4 is capable of driving IgM expression and is amenable to therapeutic targeting through BET inhibition. These data therefore highlight a novel genetic basis for ABC-like DLBCL with potential implications for future clinical studies.

## RESULTS

### DNA copy number gains of chromosome 18 are the most frequent genetic alteration in DLBCL

In order to identify significant CNAs in DLBCL, we interrogated the genomic profiles of 1,000 DLBCLs using the GISTIC2 algorithm^14^. These included high-resolution SNP microarrays from 860 previously published cases, in addition to next generation sequencing (NGS)-derived profiles from our own cohort of 140 cases (Table S1-2). Our analysis revealed 20 significant DNA copy number gains and 21 significant DNA copy number losses (false discovery rate [FDR] <0.1; Fig. 1A and Table S3). Using a subset of 448 cases for which COO subtype data was available, we identified 9 DNA copy number alterations that were significantly more frequent in ABC-like DLBCL and 11 that were significantly more frequent in GCB-DLBCL (Fisher Q-value<0.1; Fig. 1B and S1; Table S4). The most frequent genetic alteration in ABC-like DLBCL was gain of 18q21.2, which was observed in 44% of tumors. In line with the association with ABC-like DLBCL, 18q21.2 gains were associated with significantly reduced overall survival in both CHOP and R-CHOP treated patients (Fig 1C-D). Using 199 tumors with matched COO subtype, DNA copy number data and mutation status for 40 genes, we observed that the frequency of 18q21 gains (23.1% of all tumors; 40.7% of ABC-like tumors) was higher than other ABC-like DLBCL-associated somatic mutations including *MYD88* mutation (16.6% of all tumors; 33.3% of ABC-like tumors), *CD79B* mutation (7.5% of all tumors; 18.5% of ABC-like tumors) and other ABC-associated genes (Fig 1E, S2 and Table S5). Because multiple genetic alterations are associated with ABC-like DLBCL, we employed the REVEALER algorithm^15^ to identify the set of genetic alterations that best explained the ABC-like DLBCL signature. Using a set of 87 DNA copy number alterations and recurrently mutated genes as the feature set and *MYD88* mutations as the seed feature, REVEALER identified an additional 4 genetic alterations including 18q21.2 gain as those best associating with the ABC-like signature (Fig. 1F). Gains of 18q21 are therefore the most frequent genetic feature of ABC-like DLBCL and are predicted to contribute to this molecular phenotype.

**Figure 1:**
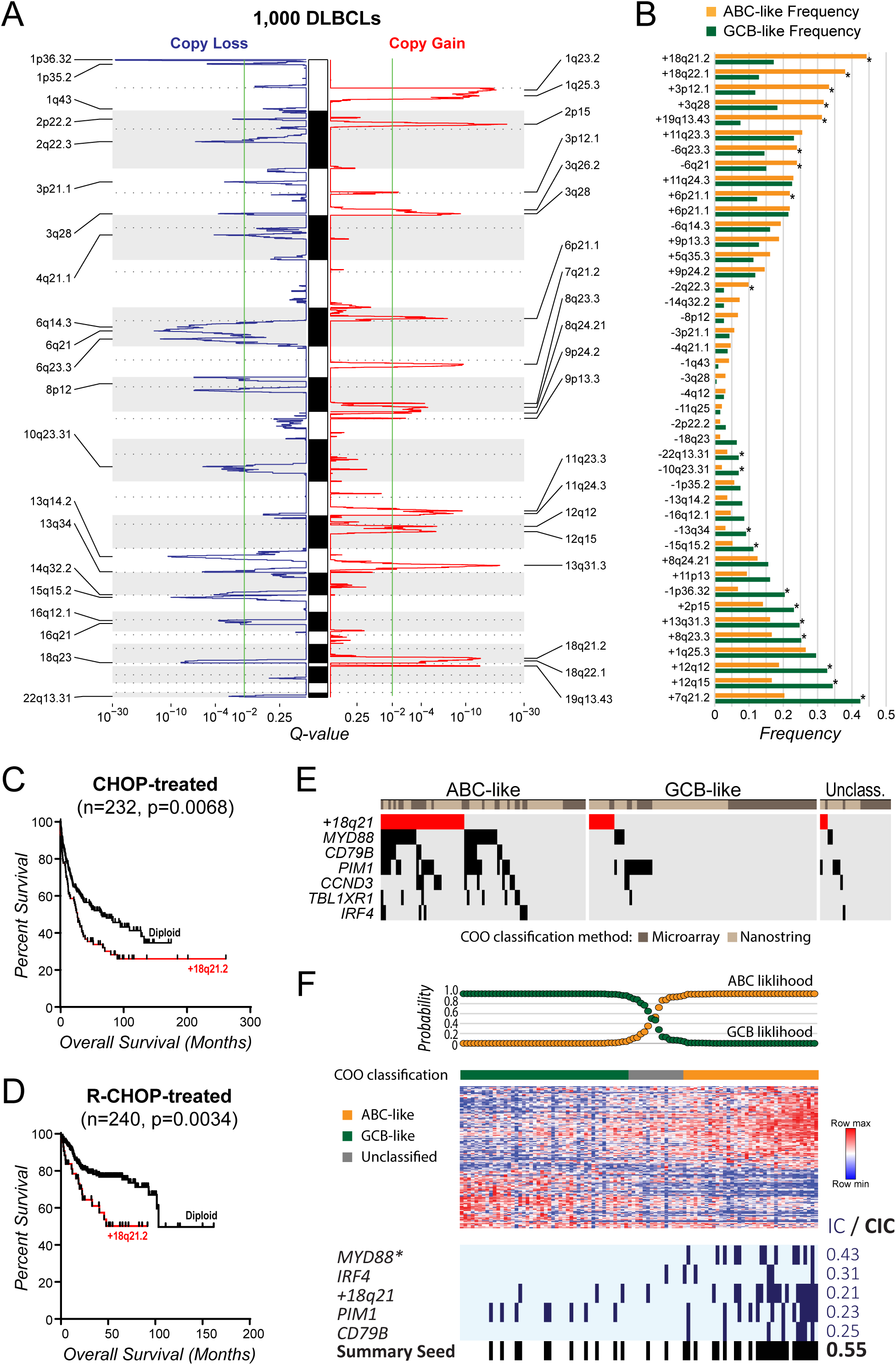
DNA copy number gains of 18q21.2 are the most frequent genetic alteration in ABC-like DLBCL. **A)** GISTIC analysis of DNA copy number profiles form 1,000 DLBCL tumors identified 21 peaks of DNA copy loss (blue, left) and 20 peaks of DNA copy gain (red, right). The green line indicates the significance threshold of q-value = 0.1. **B)** The GISTIC peaks from (A) are shown with reference to their frequency in ABC-like (orange) compared to GCB-like (green) cell of origin subtypes (*Q-value<0.1). DNA copy gains of 18q21.2 were the most frequent alteration in ABC-like DLBCL cases. **C-D)** A Kaplan-Meier plot of overall survival for patients treated with CHOP combination chemotherapy (C) or CHOP plus Rituximab (D) shows that the presence of 18q21.2 gain is associated with poor outcome. **E)** The frequency of 18q21 gains is shown relative to other somatic mutations that are significantly associated with the ABC-like DLBCL subtype. This shows that gains of 18q21 are the most frequent genetic alteration in ABC-like DLBCL. **F)** REVEALER analysis was performed to identify the set functionally-complementary genetic features that likely contribute to the ABC-like DLBCL molecular phenotype. Mutations of *MYD88* were used as the seed feature. Mutations of *IRF4, PIM1* and *CD79B*, and DNA copy gains of 18q21 were selected as additional features that likely also contribute to the phenotype (*Seed feature; IC, information coefficient; CIC, conditional information coefficient).

### The TCF4 (E2-2) transcription factor is the target of 18q21 gains in ABC-like DLBCL

Gains of 18q have been previously attributed to the *BCL2* oncogene^16,17^. However, our analysis of this large cohort provided the resolution to identify two significant peaks of DNA copy gain on chromosome 18; 18q21.2 (16 genes, Q=4.8×10^−14^) and 18q22.1 (70 genes, Q=1.1×10^−7^; Table S3). We further integrated GEP data from 249 tumors to identify the likely targets of these lesions by testing for the increase in expression of genes within the most significant peaks of DNA copy gain. This highlighted *TCF4* and *BCL2* as likely targets of the 18q21.2 and 18q22.1 gains, respectively (Fig. 2A; Table S6). Notably, most 18q copy number alterations incorporated both of these genes (Fig 2A-C). Only 7.3% or 1.0% of ABC-like DLBCLs have copy number alterations targeting *TCF4* or *BCL2* alone, respectively (Fig. 2B-C). In addition, we observed that *TCF4* was more highly expressed in ABC-like DLBCL compared to GCB-like DLBCL generally, but was further increased by DNA copy gain (Fig 2D). In line with this, ABC-like DLBCL cell lines expressed TCF4 protein irrespective of DNA copy number, but these levels were significantly increased by DNA copy gain (Fig 2E).

**Figure 2:**
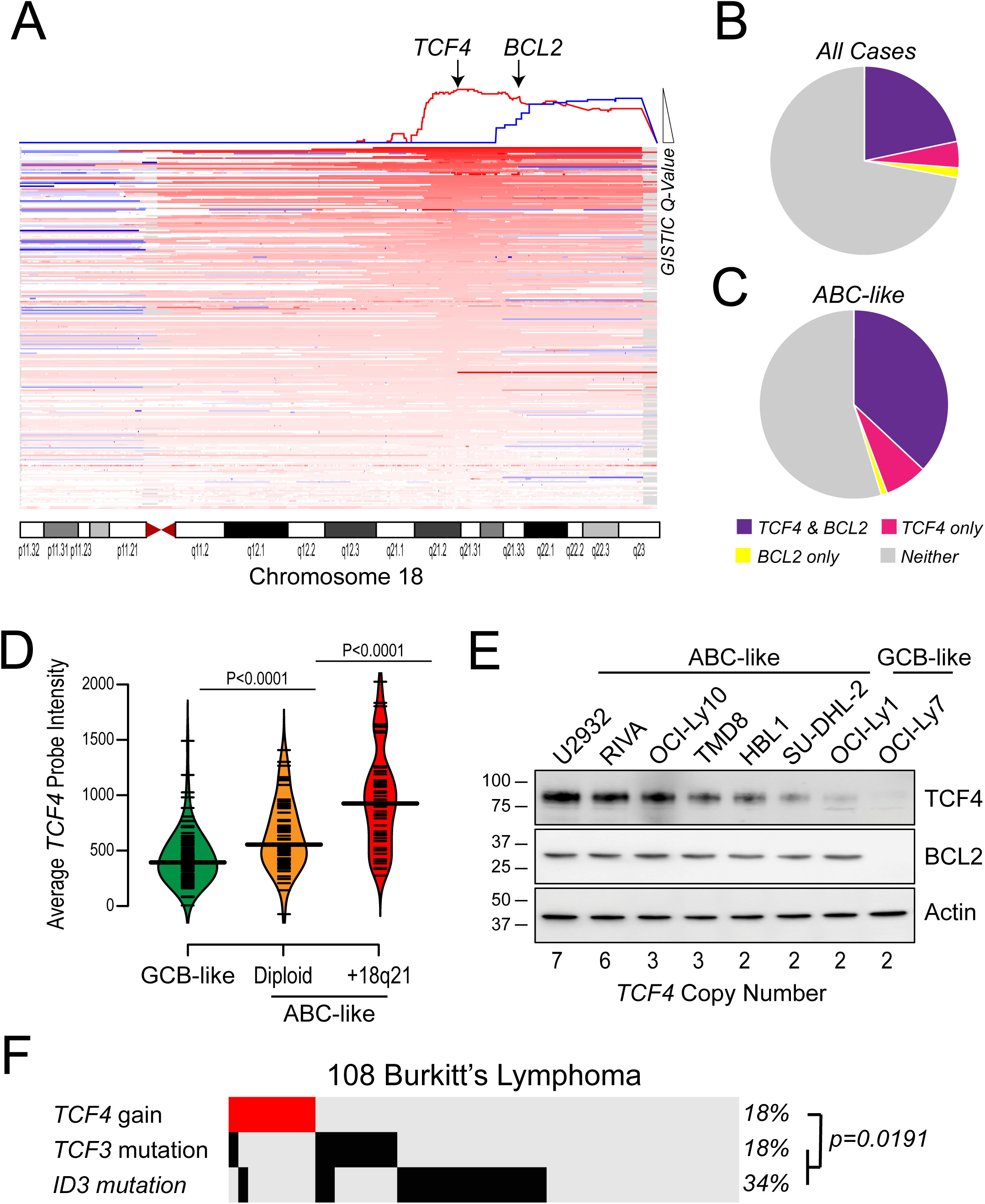
The TCF4 gene is the most significant target of 18q DNA copy number gains. **A)** A schematic of 18q DNA copy number gains is shown, with each line representing a single tumor and deeper shades of red indicating higher DNA copy number. The GISTIC q-value is shown at the top of the diagram and the two significant peaks are highlighted with arrows. The most statistically significant peak harbors the *TCF4* gene, while the less significant peak harbors the *BCL2* gene. However, it can be seen that in many cases the DNA copy number gains span both the *TCF4* and *BCL2* regions. **B-C)** The frequency of tumors with DNA copy number gains that include both the *TCF4* and *BCL2* genes (purple), the *TCF4* gene and not the *BCL2* gene (pink) or the *BCL2* gene and not the *TCF4* gene (yellow) are shown for all tumors (B) and for the ABC-like only (C). This shows that the majority of 18q DNA copy gains include both TCF4 and BCL2, but *TCF4* is more frequently gained independently of *BCL2* than vice versa. **D)** The gene expression level from microarrays are shown for GCB-like DLBCL (green) and ABC-like (orange) DLBCL tumors that are diploid for 18q, and for ABC-like DLBCL tumors that harbor *TCF4* DNA copy number gains (red). The expression of TCF4 is significantly higher in ABC-like DLBCL compared to GCB-like DLBCL in diploid cases and further significantly increased by DNA copy number gain. P-values are from students T-test. **E)** The protein level of TCF4 and BCL2 are shown in ABC-like DLBCL cell lines, ordered according to DNA copy number of the *TCF4* locus. Two GCB-like DLBCL cell lines are shown for reference. The ABC-like DLBCL cell lines express higher levels of TCF4 than the GCB-like cell lines and there is a visible relationship between TCF4 protein abundance and DNA copy number that is less clear for BCL2. **F)** The frequency of *TCF4* DNA copy gains, *TCF3* mutation and *ID3* mutation are shown for a cohort of 108 Burkitt lymphoma tumors. Gains of the *TCF4* locus are present at the same frequency of *TCF3* mutations and are significantly mutually exclusive from *TCF3* and *ID3* mutations.

The *TCF4* gene encodes an E2 family transcription factor, E2-2. Mutations of another E2 transcription factor, *TCF3*, and its negative regulators *ID2* and *ID3* are frequent in Burkitt’s lymphoma (BL) and promote immunoglobulin signaling^18,19^. We therefore interrogated the mutation status of these genes and *TCF4* copy gains in our cohort of 140 DLBCLs and a prior cohort of 108 BLs that were sequenced and analyzed with the same approach^20^. We did not observe recurrent mutations of *TCF4* or *ID2* in this BL cohort, and mutations of *TCF3* and *ID3* were infrequent in DLBCL (data not shown). However, in BL, gains of *TCF4* were present at the same frequency as *TCF3* mutations (18%). Furthermore, *TCF4* gains were significantly mutually-exclusive from *TCF3* and *ID3* mutations (Fisher P=0.019; Fig. 2F), suggesting that *TCF4* gains may serve a similar function as *TCF3*/*ID3* mutations in promoting immunoglobulin signaling. These data therefore show that the *TCF4* gene is highly expressed in ABC-like DLBCL, with expression further promoted by frequent 18q21.2 DNA copy gains, and implicates *TCF4* in immunoglobulin signaling.

### TCF4 regulates *IgM* and *MYC* expression in ABC-like DLBCL

To identify potential target genes of TCF4, we performed differential gene expression analysis of primary DLBCL tumors with TCF4 DNA copy gain (n=51) compared to those without (n=59). This analysis was limited to ABC-like tumors so as to eliminate the confounding effect of genes that differ in expression between COO subtypes. A total of 355 genes (472 probe-sets) and 87 genes (107 probe-sets) were found to be expressed at significantly higher or lower levels in tumors with *TCF4* gain, respectively (Q<0.1, fold-change≥1.2; Fig 3A; Table S7). We performed ChIP-seq of ABC-like DLBCL cell lines, SUDHL2 and TMD8, with tetracycline-inducible Myc-DDK-tagged *TCF4* in order to define whether these genes were direct transcriptional targets of TCF4 (Fig 3A). Importantly, *TCF4* was expressed at a level comparable to that in the U2932 cell line with TCF4 copy gain (Fig. S3). Using the intersection of significant peaks from both cell lines, we identified TCF4 binding proximal to 180/355 genes with increased expression and 46/87 genes with decreased expression in tumors with TCF4 copy gain (Fig. 3A-B; Table S8). These peaks showed a highly significant enrichment of motifs containing E-box consensus sequences (CANNTG; Fig. S4), and many of the same regions are also bound by TCF4 in plasmacytoid dendritic cell neoplasms^21^ (Fig. S4), providing strong evidence that we detected on-target binding. Among the most significant ChIP-seq peaks were those within the immunoglobulin heavy chain locus (Fig. 3B), in line with the significantly higher expression of *IGHM* in ABC-like DLBCL tumors with *TCF4* copy gain (Fig. 3C). This included peaks immediately upstream and downstream of the *IGHM* and *IGHD* genes, respectively, in regions with corresponding H3K27Ac in normal CD20+ B-cells that indicates they are *bona fide* enhancers (Fig. 3D). Tetracycline-inducible expression of *TCF4* in ABC-like DLBCL cell lines led to a marked increase in *IGHM* at the transcript (Fig. 3E) and protein level (Fig. 3F). In comparison, BCL2 expression was not induced by TCF4 over-expression and MYC induction was restricted to the two cell lines that lacked *MYC* translocation (SUDHL2 and TMD8; Fig. S5). These data show that IgM is a direct target of TCF4 and can be induced by its over-expression in ABC-like DLBCL.

**Figure 3:**
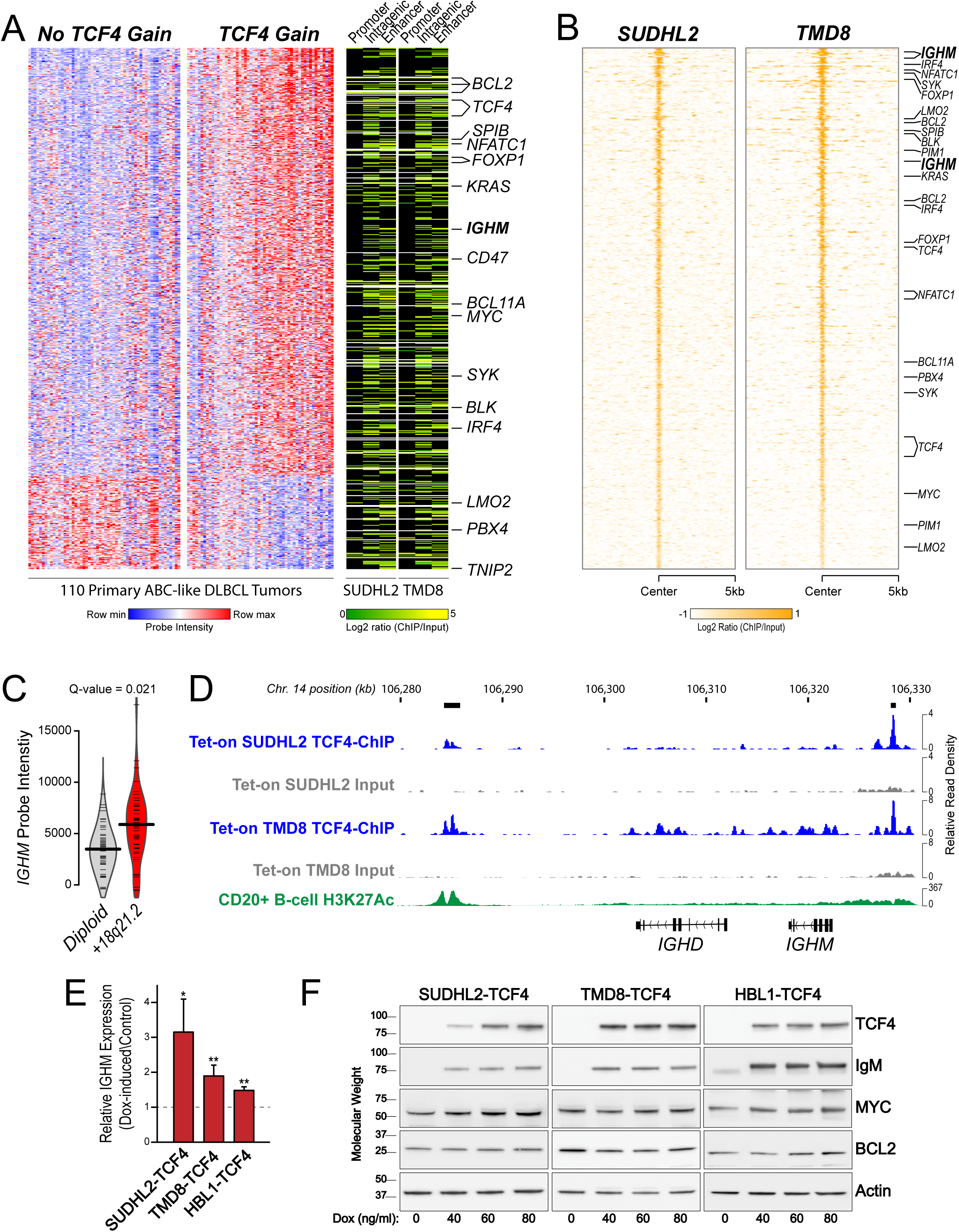
TCF4 regulates IgM expression in ABC-like DLBCL. **A)** Differential gene expression analysis of 110 primary ABC-like DLBCL tumors with or without *TCF4* DNA copy number gain identified a large set of genes with increased expression associated with *TCF4* gain. This included the direct targets of 18q DNA copy number gains, *TCF4* and *BCL2*, and multiple genes with an important role in the pathophysiology of DLBCL, such as *IRF4, MYC* and the immunoglobulin heavy chain μ (*IGHM*), that are upregulated as a secondary effect of *TCF4* gain. ChIP-seq of TCF4 from SUDHL2 and TMD8 cell lines showed that the majority of these genes were marked with *TCF4* binding in intragenic or distant enhancer elements, suggesting that their up-regulation may be driven by transcriptional activation by TCF4. **B)** The significant TCF4 ChIP-seq peaks from SUDHL2 and TMD8 are shown, ordered from strongest (top) to weakest (bottom) signal ratio compared to the input control. Significant peaks were detected for important genes such as *MYC* and *IRF4*, but multiple *IGHM* peaks were amongst those with the highest TCF4 binding. **C)** A violin plot shows that primary DLBCL tumors with 18q21 gain express significantly higher transcript levels of IGHM. **D)** Two of the TCF4 peaks at the immunoglobulin heavy chain locus are shown for TCF4 ChIP (blue) compared to the equivalent input control (grey). A black box indicates the significant peak. For reference, ENCODE data for H3K27 acetylation (H3K27Ac) ChIP-seq in CD20+ B-cells is shown, which support the TCF4 bound regions as bona fide enhancer elements in B-cells. **E)** Tetracycline-induced expression of TCF4 in ABC-like DLBCL cell lines with low TCF4 copy number resulted in a significant increase in IGHM transcript compared to control cells. **F)** Tetracycline-induced expression of TCF4 led to a marked increase in IgM protein in ABC-like DLBCL cell lines with low TCF4 copy number. An increase in MYC was also observed in SUDHL2 and HBL1 cell line, but was not significant in TMD8. No change was observed for BCL2. The quantification of triplicate experiments is shown in Figure S5.

### TCF4 can be targeted by the BET proteolysis-targeting chimera (PROTAC), ARV771

The *TCF4* gene is one of the most highly BRD4-loaded genes in DLBCL, including in ABC-like DLBCL cell lines with *TCF4* copy gain (Fig. S6). We therefore evaluated small molecule BET inhibitors and a BET protein degrader, ARV771, as a potential avenue for reducing TCF4 expression in ABC-like DLBCL cell lines with high-copy number of *TCF4*. The small molecule BET inhibitors, JQ1 and OTX015, resulted in an up-regulation of BRD4 that was not observed with ARV771 due to its role as a sub-stoichiometric BRD4 degrader (Fig 4A and S7). This was associated with a greater efficacy of ARV771 in reducing the BRD4 target genes, MYC and TCF4 (Fig 4A), and the ability of ARV771 to induce apoptosis of these cell lines (Fig 4B). However, as MYC is also a target of TCF4, the down-regulation of MYC is likely partially mediated through TCF4.

**Figure 4:**
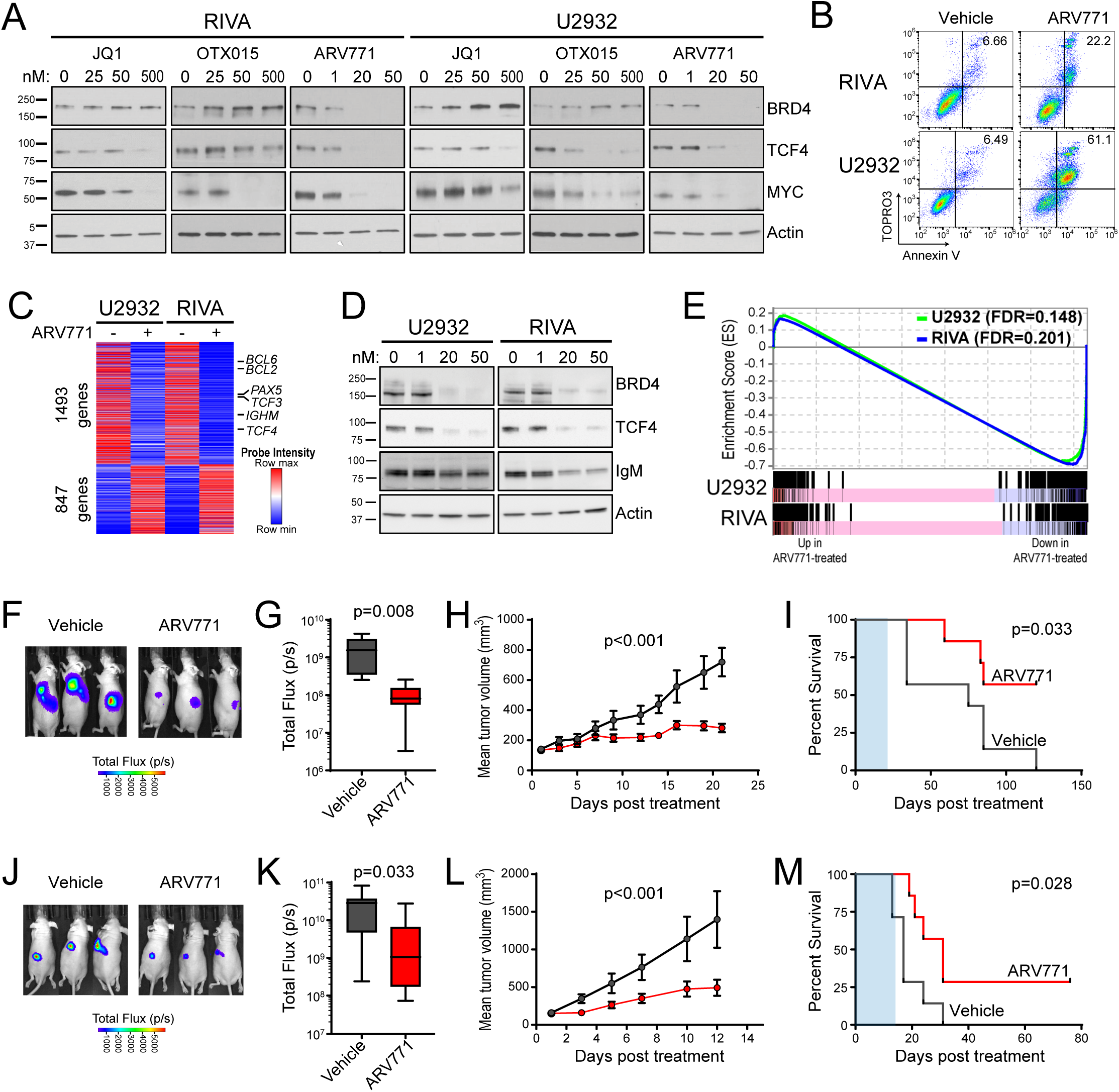
BET proteolysis-targeting chimeras (PROTAC) effectively inhibit TCF4 and show efficacy in ABC-like DLBCL cell lines. **A)** The treatment of ABC-like DLBCL cell lines with high *TCF4* DNA copy number using small molecule BET inhibitors, JQ1 and OTX015, leads to an accumulation of BRD4 but can reduce the BRD4-targets TCF4 and MYC. The BET PROTAC, ARV771, effectively degrades BRD4 and leads to a more potent reduction of TCF4 and MYC at 10-fold lower doses than the small molecule inhibitors. **B)** Treatment of U2932 and RIVA cell lines with 50nM of ARV771 for 48h led to the induction of apoptosis, as measured by TOPRO3 / Annexin-V positive staining. **C)** The treatment of U2932 and RIVA cell lines with 50nM of ARV771 for 24h led to broad changes in transcript levels. This included the down-regulation of known BRD4 target genes, *BCL6* and *PAX5*, as well as the down-regulation of *TCF4* and it’s target gene, *IGHM*. **D)** The down-regulation of IgM following treatment with ARV771 is also observed at the protein level. The quantification of triplicate experiments is shown in Figure S7. **E)** Gene set enrichment analyses are overlaid for U2932 (green) and RIVA (blue) for the set of genes that were more highly expressed in primary ABC-like tumors with TCF4 DNA copy number gain compared to those tumors without DNA copy number gain, as shown in Figure 3A. Treatment with ARV771 led to a significant and coordinate down-regulation of this TCF4-associated signature in both the U2932 and RIVA cell lines. **F-I)** Murine xenografts of the U2932 cell line were allowed to become established and then treated with 30mg/kg of ARV771 daily x 5 days per week for 3 weeks. At the end of treatment the luminescence was significantly lower in ARV771-treated mice compared to vehicle control (F, G) as a result of the significant reduction in tumor growth (H). This led to significantly prolonged survival in ARV771-treated mice. **J-M)** Murine xenografts of the RIVA cell line were allowed to become established and then treated with 30mg/kg of ARV771 daily x 5 days per week for 2 weeks. At the end of treatment the luminescence was significantly lower in ARV771-treated mice compared to the vehicle control (J-K), as a result of the significant reduction in tumor growth (L). Despite the short duration of treatment, this led to a significant prolongation in survival of ARV771-treated mice (M).

Reductions of TCF4 by ARV771 treatment were accompanied by reduced expression of the TCF4 target genes. This included significant reductions of IgM at the transcript and protein level (Fig. 4C-D and Table S9). ARV771 treatment led to significant down-regulation of the set of genes that were identified as being increased in association with *TCF4* DNA copy number gain in primary tumors (Fig. 4E). The promising *in vitro* activity of ARV771 led us to test whether this compound would be efficacious *in vivo*. In xenografts of the U2932 (Fig. 4F-I) and RIVA (Fig. 4J-M) cell lines that express high levels of TCF4, we observed that ARV771 was able to significantly reduce tumor growth. At the end of treatment, tumors were significantly smaller in ARV771-treated mice and this led to a significant prolongation of survival in these mice (Log Rank P-value < 0.05). Together these data demonstrate a clear functional rationale for BET inhibition in ABC-like DLBCL, and show that ARV771 is effective at eliminating TCF4 and its target genes and treating ABC-like DLBCL *in vivo*.

## DISCUSSION

The ABC-like subtype is one of two major molecular subtypes of DLBCL that is recognized by the WHO classification^22^. These tumors are driven by chronic active B-cell receptor signaling that emanates from autoreactive IgM that is localized to the cell surface and intracellular lysosomes^8,23-25^. Mutations in *CD79B* and *MYD88* deregulate this signaling through the reduction of LYN-mediated negative feedback and by activation of IRAK signaling, respectively^8,10^. However, recent murine studies have shown that *MYD88* mutation alone drove a phenotype that was reminiscent of peripheral tolerance, and this was only relieved by the combination of *MYD88* and *CD79B* mutations together, or by increased expression of surface IgM^26^. The ABC-like phenotype is therefore the result of cumulative epistatic genetic alterations, rather than a single dominant driver mutation. In further support of this notion, recent genomic studies have defined co-associated sets of genetic alterations that co-segregate with unique genetic subsets of ABC-like DLBCL^27^. The “cluster 5” subset of ABC-like DLBCL included frequent *MYD88* and *CD79B* mutations, but the most frequent genetic alteration in this subtype was DNA copy number gain of 18q^27^.

We identified the *TCF4* (aka E2-2) gene as the most significant target of 18q DNA copy number gains in DLBCL. The *TCF4* gene (aka E2-2) is closely related to *TCF3* (aka E2A), with both encoding helix-loop-helix transcription factors that form dimers and recognize E-box consensus sequences (CANNTG)^28^. Murine conditional knock-out studies showed that TCF3 and TCF4 are critical regulators of germinal center B-cell and plasma cell development, in part due to their role in activating immunoglobulin heavy- and light-chain enhancer elements^29,30^. The ID2 and ID3 proteins bind to and inhibit the activity of TCF3 and TCF4 by preventing their dimerization and DNA binding^28^. The *TCF3* and *ID3* genes are recurrently mutated in another form of B-cell lymphoma, Burkitt Lymphoma, with the mutations residing in the interface between *TCF3* and *ID3* and preventing their interaction^18,19^. Mutations in *ID3* are approximately twice as frequent as mutations of *TCF3*, and presumably also reduce the interaction between ID3 and TCF4 considering the high degree of homology between these two proteins. We observed that TCF4 DNA copy number gains are also frequent in Burkitt lymphoma, and that they mutually exclude *TCF3* and *ID3* mutations, providing further evidence for the importance of *TCF3*/*TCF4* deregulation in this disease. In contrast to Burkitt lymphoma, *TCF3* and *ID3* mutations are rare in ABC-like DLBCL, but *TCF4* DNA copy number gains are present at more than twice the frequency. In line with the murine studies, we observed a marked up-regulation of IgM transcript expression in primary tumors with *TCF4* DNA copy number gain. We also identified binding sites for TCF4 in the immunoglobulin heavy chain locus and showed that induced expression of TCF4 was sufficient for increasing the expression of IgM at the transcript and protein level. Together, this provides strong evidence for a functional role of TCF4 in promoting IgM expression in ABC-like DLBCL. This is particularly important in this disease, because >90% of ABC-like DLBCL cases express IgM and the disease etiology centers on pathogenic signaling downstream of this receptor^7,8^. Notably, *TCF4* was more highly expressed in ABC-like DLBCL compared to GCB-like DLBCL generally, even in cases without DNA copy number gain of the locus. This suggests that this axis may be active in all ABC-like DLBCLs, and further enhanced in the ∼40% that harbor 18q DNA copy number gains. This is akin to the role of EZH2 in GCB-like DLBCL, which promotes the survival and proliferation of all germinal center B-cells but has enhanced activity in the context of hypomorphic somatic mutations^31,32^. We therefore hypothesize that *TCF4* may participate in a critical functional axis of immunoglobulin regulation in all ABC-like DLBCL.

Proteins in the BET family, including BRD4, are attractive therapeutic targets in cancer due to their role in the transcriptional activation of oncogenes such as *MYC*^33,34^. In DLBCL, BRD4 targets include key transcription factors such as *BCL6, PAX5* and *IRF4*^35^. We have highlighted *TCF4* as another prominent target of BRD4 in DLBCL, as has been previously described in plasmacytoid dendritic cell neoplasms^21^. Due to the difficulty in directly drugging transcription factors, BET inhibition therefore represents a logical avenue for reducing TCF4 expression in ABC-like DLBCL. Notably, cell line studies have shown that the small molecule BET inhibitor OTX015 induces apoptosis in ABC-like DLBCL cell lines, as compared to a predominantly cytostatic effect in GCB-like DLBCL cell lines^36^. However, small molecule inhibitors have also been shown to result in the up-regulation of BRD4 expression^37^. We therefore evaluated a novel BET protein PROTAC, ARV-771, which combines a BET-targeting warhead from OTX015 with a moiety that recruits the VHL ubiquitin ligase^37^. Because the PROTAC is not degraded, this results in sub-stoichiometric proteolysis of BET proteins, including BRD4. We found that ARV-771 was able to inhibit the expression of TCF4 at 10-fold lower concentrations than small molecule BET inhibitors. The inhibition of TCF4 expression in ABC-like DLBCL cell lines with high TCF4 copy number was accompanied by the coordinate down-regulation of genes that were highly expressed in primary tumors with TCF4 DNA copy gain, suggesting that a subset of the broad transcriptional changes associated with BRD4 degradation were the consequence of reduced TCF4. This was also associated with the induction of apoptosis at low nanomolar doses of ARV-771, significant reduction of tumor growth *in vivo*, and a significant prolongation in the life of tumor-bearing animals. Together, this highlights *TCF4* DNA copy gains as a functional rationale for BET inhibition in ABC-like DLBCL and shows that the BET PROTAC ARV-771 has significant activity in this context. The over-expression of BCL2 has been described as a resistance mechanism for BET inhibitors^38^. We observed that the majority of 18q DNA copy number gains in DLBCL encompass both the *TCF4* and the *BCL2* gene, and we therefore posit that the promising activity of BET inhibitors in ABC-like DLBCL may be further enhanced by combination with a BCL2 inhibitor such as Venetoclax. In support of this, BET inhibitors have been shown to act synergistically with Venetoclax in myeloid leukemia^38^ and in another form of B-cell lymphoma, mantle cell lymphoma^39^. Combination of BET and BCL2 inhibition therefore represents an attractive therapeutic avenue for future investigation in ABC-like DLBCL.

In conclusions, we have identified DNA copy number gains of TCF4 as the most frequent genetic alteration in ABC-like DLBCL. Increased expression of TCF4 leads to its occupancy on IgM enhancer elements and increased expression of IgM at the transcript and protein level. We have shown that BET-targeting PROTACs efficiently reduce the expression of TCF4 and its target genes, induce apoptosis in ABC-like DLBCL cells, and prolong the life of mice bearing ABC-like DLBCL tumors. This study therefore highlights the BRD4-regulated TCF4 and IgM axis as a functional rational for the use of BET inhibitors in ABC-like DLBCL.

## MATERIALS AND METHODS

### DNA copy number data acquisition and processing

Publicly available data for single nucleotide polymorphism microarrays and array comparative genome hybridization platforms with >200,000 markers were downloaded from the gene expression omnibus^16,17,40-46^ (Table S1; www.ncbi.nlm.nih.gov/geo/). These included Affymetrix 250K and SNP 6.0 platforms, and the Agilent 244K platform. For Affymetrix microarrays, raw CEL files were extracted and copy number predicted using the Affymetrix Copy Number Analysis for Genechip (CNAG) tool, with reference to data from 100 Caucasian HapMap samples. Agilent data was analyzed using BioConductor, as previously described^17^. Data for all arrays were represented as Log2 copy number change and segmented using the circular binary segmentation (CBS) tool in GenePattern^47^. Peaks of significant DNA copy number loss and gain were identified using GISTIC2.0^14^. The thresholds utilized for DNA copy number gain and loss were 0.2 copies over a region encompassing 100 markers.

### Targeted next generation sequencing (NGS) and variant calling

Genomic DNA for 140 fresh/frozen DLBCL tumors were obtained from the University of Nebraska Medical Center (UNMC) lymphoma tissue bank (IRB 161-95-EP) and interrogated by targeted sequencing of a panel of 380 genes, as previously described^20^. In brief, 500-1,000ng of genomic DNA was sheared using a Covaris S2 instrument and libraries prepared using KAPA Hyper Prep kits (Roche) and Illumina TruSeq Adapters (Bioo Scientific) according to the manufacturer’s protocol. A maximum of 6 cycles of PCR was used for library preparation. Samples were 12-plexed, subjected to hybrid capture with a 5.3Mbp Nimblegen SeqCap custom reagent (Roche), and amplified by 8 cycles of PCR. Each pool was sequenced on a single lane of an Illumina HiSeq 2500 instrument in high-output mode using 100bp paired-end reads at the Hudson Alpha Institute for Biotechnology Genome Sequencing Laboratory. Raw sequencing reads were aligned to the human genome (hg19) using BWA-Mem^48^, realigned around InDels using GATK^49^, sorted and deduplicated using Picard tools, and variants were called according to a consensus between VarScan2^50^ and GATK Unified Genotyper^49^. This approach has been validated to have a specificity of 92.9% and a sensitivity of 86.7%^51^. Average on-target rate for this dataset was 88% and average depth of coverage 623X (min = 122X, max = 1396X). Raw FASTQ files for the targeted NGS of previously published DLBCLs (n=119; European Nucleotide Archive Accession ERP021212)^52^ and Burkitt’s lymphomas^20^ were also analyzed with the same pipeline, and the results integrated. The DNA copy number of UNMC DLBCL and Burkitt lymphoma cohorts was determined using CopyWriteR^53^ with 200kB windows.

### DLBCL Cell Lines

The SU-DHL-2 cell line was obtained from ATCC. The RIVA (aka RI-1), HBL1, TMD8, U2932 and OCI-Ly10 cell lines were obtained from DSMZ. The DNA copy number profile of DLBCL cell lines were derived from previously reported SNP6.0 data^54^ or targeted next generation sequencing, as described above. Cell of origin subtype was determined according to previous descriptions^8^. U2932, RIVA, TMD8, HBL1, and SUDHL2 were maintained in RPMI-1640 media with 10% FBS and 1% penicillin/streptomycin. OCI-LY1, OCI-LY7 and OCI-LY10 were maintained in IMDM supplemented with 20% human serum and 1% penicillin/streptomycin. Cell lines were regularly tested for mycoplasma, and identity confirmation by Short Tandem Repeat at core facility of MD Anderson Cancer Center. Tetracyline-inducible expression of *TCF4* was performed in the TMD8, HBL1, and SUDHL2 cell lines. Detailed methodology can be found in the supplementary methods.

### ChIP-sequencing of TCF4

For inducible TCF4 expression, TMD8-TCF4, or SU-DHL-2-TCF4 cell lines were treated with doxycycline (60ng/ml) for 24 hours. For chromatin immuno-precipitation, five million cells were fixed with 1% formaldehyde for 10 min, quenched by addition of 125 mM Glycine for 5 min at RT, washed with ice-cold PBS then resuspended and incubated in ice-cold ChIP buffer (10mM Tris-HCl pH 8.0, 6.0 mM EDTA, 0.5% SDS and protease inhibitor) for 1hour. In the same time, 5μg of antibodies (TCF4^21^) or control rabbit IgG (Cell Signaling; 2729) were allowed to bind to dynabeads Protein-G (Invitrogen; 10003D) in binding buffer (0.2% BSA, 0.1% Tween-20 in PBS) for 2 hours. Chromatin was sheared using Covaris M220. Sonicated lysates were diluted in dilution buffer (10mM Tris-HCl pH8, 140 mM NaCl, 1mM EDTA pH 8, 0.5 mM EGTA, 1% Triton X-100, and 0.1% Sodium Deoxycholate) and added to antibody bound Protein-G beads for immunoprecipitation overnight at 4ºC. Note; for ChIP normalization, spike in chromatin/antibody was added to sonicated lysates (53083/61686; Active Motif). Next day, bead-bound complexes were washed 5 times with RIPA buffer (1% NP40, 0.1% SDS, and 0.5% Sodium Deoxycholate in PBS), 2 times with LiCl buffer (10 mM Tris-HCl pH8, 250 mM LiCl, 0.5% NP40, 0.5% Sodium Deoxycholate, 1 mM EDTA), once with TE buffer pH 8.0 and finally resuspended in 50μl of TE buffer containing 20μg of proteinase K and RNase A (0.2 μg/μl). TE buffer, RNase A and proteinase K mixture was also added in total chromatin samples in parallel as input reference. Reverse cross-linking was performed at thermal cycler (4 hours 37^°^C, 4 hours 50^°^C, and overnight 65^°^C). DNA purification was performed with SPRIselect beads (Beckman Coulter; B23317) and further processed for library generation with KAPA HyperPrep kit (KK8502) according to the kit protocol.

Sequencing reads were aligned to the human genome (hg19) using BWA-Mem^48^, realigned around InDels using GATK^49^, sorted and deduplicated using Picard tools. Peaks were called in TCF4 ChIP samples compared to their input control using EaSeq with global thresholding. Peaks were annotated according to the transcription start site of the nearest RefSeq gene and filtered based upon FDR (<0.1), log2ratio of TCF4 ChIP vs. isotype control (≥2.0), peaks that overlapped between TMD8 and SUDHL2, and peaks corresponding to genes with differential expression between ABC-like DLBCL tumors with or without TCF4 DNA copy number gain. Peaks within 2kbp of the transcription start site were defined as ‘promoter’ peaks, those outside of the promoter region but within the coding region of the gene were defined as ‘intragenic’ peaks, and those outside of these regions but within 50kbp of the transcription start site were defined as distant ‘enhancer’ peaks. For visualization, files were converted to wiggle format and viewed using the Integrative Genomics Viewer^55^. The wiggle file for H3K27Ac ChIP-seq for CD20+ B-cells was downloaded directly from UCSC Genome Browser (https://genome.ucsc.edu/ENCODE/downloads.html). Significantly over-represented DNA sequence motifs (FDR<0.05) were identified in TCF4 ChIP-seq peaks compared to the reference genome (hg19) using CisFinder^56^ with the default settings. Motifs with 75% homology were collapsed to motif clusters.

For ChIP-PCR, chromatin immuno-precipitation was performed with BRD4 antibody (Bethyl, Cat No. A301-985A) following the protocol as described above for TCF4. Chromatin DNA was also purified from the input samples. The purified DNA was used to perform quantitative PCR using SYBR Green/ROX qPCR Master Mix (Applied Biosystem; 4309155). Percentage of input was quantified from the adjusted input Ct values and further used to determine ΔCt values for BRD4 or IgG ChIP. Primers used for ChIP-PCR have been listed in Table S10.

### BET Inhibitors and Treatments

BRD4 inhibitors JQ1 and OTX015 were obtained from Selleck Chemicals. BRD4-PROTAC (ARV-771) was provided by Arvinas, Inc. (New Haven, CT). U2932 and RIVA cell lines were treated with indicated concentrations of BET-inhibitors (JQ1, OTX015) or BRD4-PROTAC (ARV771) for 24 hours before immunoblotting. For apoptosis analysis, U2932 and RIVA cell lines were seeded at 2.5 × 10^5^ cells/ml and treated with ARV771 at indicated concentration for 48 hours. Cells were stained with Annexin V (Thermo Fisher; A35122)/To-PRO-3 and analyzed using flow cytometry (BD LSRFortessa) and FlowJo software. For gene expression analysis, cell lines (U2932 and RIVA) were un-treated or treated with ARV771 (50ng/ml) for 24 hours. Total RNA was extracted using All prep DNA/RNA kit (Qiagen; 80204) and RNA integrity was assessed using an Agilent-4200 TapeStation system. Libraries were generated using KAPA RNA HyperPrep kit with RiboErase (KK8560) according to the manufacturer’s instructions. Libraries were pooled and run on a single land of a HiSeq 4000 instrument at the MD Anderson Sequencing and Microarray Core Facility. Fastq files were first aligned to the GRCh37 assembly with GENCODE37lift37 annotations using STAR 2.6.0c, using a two-pass protocol with alignment parameters from the ENCODE long RNA-seq pipeline. The transcript-coordinate output files were then pre-processed with RSEM version 1.2.31’s convert-sam-for-rsem tool before quantifying with rsem-calculate-expression, assuming the data is from an unstranded paired-end library. Tximport version 1.6.0 was then used under R version 3.4.3 to read individual RSEM output files and aggregate to gene-level expressions based on the gene-transcript relationships in GENCODE27lift37’s Gene symbol metadata. DESeq2 version 1.18.1 was used to identify differentially expressed genes using a two-variable (Cell line and Treatment) analysis with default settings. Gene set enrichment analysis57 was performed using GenePattern and a list of all genes from RNA-seq ranked by the fold-change in expression following ARV771 treatment. The gene set consisted of all genes that showed significantly higher expression in ABC-like DLBCL tumors with *TCF4* DNA copy number gain compared to those without, as shown in Figure 3.

### Murine Xenograft Experiments

#### Reagents and antibodies

ARV-771 was kindly provided by Arvinas, Inc. (New Haven, CT) D-Luciferin (potassium salt) was obtained from Gold Biotechnology, Inc. (St Louis, MO). BD Matrigel Matrix High Concentration was obtained from BD Biosciences (Franklin Lakes, NJ) (Catalog number 354248).

#### Cell lines

Luciferase-expressing RIVA and U2932 cells were created by transducing cells with Luc-ZSGreen. pHIV-Luc-ZsGreen was a gift from Bryan Welm (Addgene plasmid # 39196). High GFP-expressing cells were isolated by flow sorting for GFP expression in the M. D. Anderson Flow Cytometry and Cellular Imaging Core Facility (FCCICF), a shared resource partially funded by NCI Cancer Center Support Grant P30CA16672.

#### In vivo studies

All animal studies were performed under a protocol approved by the IACUC at M.D. Anderson Cancer Center, an AAALAC-accredited institution. Five million RIVA or U2932 cells (mixed with Matrigel at a volume ratio of 1:1) were subcutaneously injected in the left flank of male athymic nude mice (nu/nu) (n = 8 per group). Tumor volume was calculated by the 1/2(length x width^2^) method. Treatment was initiated when the mean tumor volumes reached ∼150 mm3. Mice were treated with vehicle (10% [1:1 solutol: ethanol] and 90% D5-water, s.c. daily x 5 days per week) or ARV-771 (30 mg/kg, s.c., daily x 5 per week). The RIVA mouse model was treated for two weeks. Due to slower tumor growth, the U2932 mouse model was treated for three weeks. For bioluminescent imaging, mice were IP-injected with 100 μL of 75 mg/kg D-Luciferin potassium salt (reconstituted in 1X PBS and sterile-filtered through a 0.2 um filter) incubated for 5 minutes, anesthetized with isoflurane and imaged once per week utilizing a Xenogen IVIS-200 imaging system (PerkinElmer) to monitor disease status and treatment efficacy. One mouse from each cohort was euthanized after three weeks of treatment for biomarker analysis. Mice bearing tumors greater than 1500 mm^3^ were removed from study and humanely euthanized (carbon dioxide inhalation and cervical dislocation) according to the IACUC-approved protocol. Veterinarians and veterinary staff assisting in determining when euthanasia was required were blinded to the experimental conditions of the study. Tumor size was compared among cohorts by unpaired t-test. The survival of the mice is represented by a Kaplan Meier plot. Differences in survival were calculated by a Mantel-Cox log-rank test. P values less than 0.05 were considered significant.

## ACKNOWLEDGEMENTS

This research was supported by the Nebraska Department of Health and Human Services (LB506 2016-16, M.R.G.), the Schweitzer Family Fund (J.W.), RO1 CA210250 (K.B.) and the MD Anderson Cancer Center NCI CORE Grant (P30 CA016672). Arvinas, Inc. kindly provided ARV-771 for the studies.

## AVAILABILITY OF DATA

The data produced in this study are available in the gene-expression omnibus (www.ncbi.nlm.nih.gov/geo/), accession number GSE119241. The SNP and gene expression microarray accessions for the previously published data are listed in Table S1. Raw next generation sequencing data will be provided upon reasonable request to the corresponding author and the completion of confidentiality non-disclosure and material transfer agreements.

## AUTHOR CONTRIBUTIONS

NJ, KH, ST, WF, DK and OH performed experiments. NJ, KH, ST, KB and MRG analyzed data and wrote the manuscript. MJM, AB, TH, QD, DM, CP, AG, SR, JI, FG, SSN, JW, RED and KB analyzed or interpreted data. AA and CLL provided computational resources. EH, RK, KES, GJ, RR, RDG, AR, JV, ML, and TG provided samples or data. MRG conceived and supervised the study. All authors read and approved the manuscript.

